# The neuroeconomics of individual differences in saccadic decisions

**DOI:** 10.1101/2022.06.03.494508

**Authors:** Tobias Thomas, David Hoppe, Constantin A. Rothkopf

**Affiliations:** Centre for Cognitive Science & Institue of Psychology, Technical University Darmstadt, Germany; Hessian center for Aritifical Intelligence (hessian.ai), Darmstadt, Germany

**Keywords:** eye movements, neuroeconomics, decision-making

## Abstract

Active exploration of the visual environment requires choosing the next gaze target, which has been characterized as depending on attentional biases and on oculomotor biases. However, disentangling these two factors has provided contradictory results. Here, we conceptualize active gaze selection as a decision-making process in the context of neuroeconomics, allowing us to employ a new saccadic preference elicitation experiment and to quantify behavioral biases with a random utility model. Subjects showed similar preferences for shorter amplitude saccades in line with previous research and shared a preference for saccades back towards the previous gaze location, contrary to numerous reports on spatial “inhibition of return”. Surprisingly, the experiment revealed marked individual differences in the preferences for global gaze directions, which were stable across most participants after one week. Thus, this study provides the first measurement of intrinsic saccadic utilities independently of spatial image properties and quantitatively dissociates stable individual differences in global gaze selection.

## 1 Introduction

Humans carry out more than 100,000 eye movements every day and each one of these gaze shifts requires a decision where to look next. The reason is, that the inhomogeneous acuity across the visual field necessitates sequentially repositioning of the high acuity fovea using saccadic eye movements to acquire relevant information from the environment (Findlay & Gilchrist, 2003; Hayhoe & Ballard, 2005; Land & Tatler, 2009). As a consequence, these decisions which eye movement to carry out next fundamentally influence where we look and what visual information can be acquired. Accordingly, the active exploration of the visual environment has been characterized as depending both on attentional priority that makes certain locations within a scene more likely to be looked at (Findlay & Gilchrist, 2003; Hayhoe & Ballard, 2005; Henderson, 2003; Land & Tatler, 2009; Yarbus, 1967) and on oculomotor biases that make particular gaze shifts more likely (A. J. Anderson et al., 2008; Bays & Husain, 2012; Luke et al., 2014; Tatler & Vincent, 2009). Thus, the sequential decision process underlying gaze selection needs to integrate extrinsic visual relevance with intrinsic behavioral costs. Therefore, to understand the active visual exploration of the visual environment, it is crucial to quantify the intrinsic behavioral utilities of saccadic choices.

While visual factors determining the selection of the next gaze target have been investigated extensively (Borji et al., 2013; Henderson, 2003; Navalpakkam et al., 2010; Rothkopf et al., 2007; Schütz et al., 2012; Torralba et al., 2006), intrinsic behavioral biases, often termed oculomotor biases, have received comparatively less attention. The approaches for investigating such intrinsic behavioral biases can be divided into experimental paradigms that involve controlled laboratory settings with sparse displays allowing systematic manipulations of task variables (A. J. Anderson et al., 2008; Gilchrist & Harvey, 2000; Horowitz & Wolfe, 1998; Peterson et al., 2001) and paradigms that try to infer biases from large collections of eye movements obtained with participants looking at images of natural scenes (Bays & Husain, 2012; Luke et al., 2014; Tatler & Vincent, 2009; Wilming et al., 2013). However, the results of these investigations have revealed puzzling discrepancies and inconsistencies. For example, while some studies have found spatial biases away from recent gaze targets termed “spatial inhibition of return” (Bennett & Pratt, 2001; Klein & MacInnes, 1999), other studies have found no such bias (Horowitz & Wolfe, 1998), some studies have found the bias only in some task conditions (Dodd et al., 2009), and some studies instead found the opposite bias, “facilitation of saccadic return” (Smith & Henderson, 2009). Taken together, it is currently unknown, how to experimentally isolate behavioral biases in saccadic eye movements from attentional biases and how to quantify their contribution to the decision of the next gaze target.

One reason for the mixed results lies in the fact that spatial preferences for gaze selection interact with spatial properties of images. A clear demonstration is that the often reported oculomotor bias for horizontal versus oblique saccades (Gilchrist & Harvey, 2006) disappears by simply rotating the image being looked at (Foulsham et al., 2008). Thus, the likelihood of certain saccadic movements found in large data sets of participants viewing images cannot be attributed solely to behavioral biases. Second, experiments with sparse displays involving systematic manipulations of task variables commonly instruct subjects to saccade to a single gaze target (A. J. Anderson et al., 2008). Thus, these experimental paradigms do not involve a saccadic decision. A third reason is that while classic studies have predominantly reported average gaze behavior across many participants, more recent studies have revealed significant individual differences in oculomotor behavior (Bargary et al., 2017; Henderson & Luke, 2014; Risko et al., 2012). Nevertheless, it is unknown which factors contribute to the observed individual differences in the likelihood of certain gaze shifts and how to describe these factors computationally. Recently, intrinsic factors of visual behaviors conceptualized as behavioral costs in the framework of decision-making have been investigated more carefully (Hoppe & Rothkopf, 2016, 2019; Lisi et al., 2019; Petitet et al., 2021) but the costs of saccadic gaze shifts have been captured with generic cost functions determined by the specific task under investigation. Taken together, it is still unclear, how the cost of an eye movement influences saccadic decisions and which factors are responsible for interindividual differences of choices.

Here, we conceptualize saccadic gaze shifts as a decision making process (Gottlieb, 2012; Hayhoe & Ballard, 2014; Hoppe & Rothkopf, 2019) allowing us to adopt established and principled methods from neuroeconomics (Glimcher & Fehr, 2013; Körding et al., 2004). In this context, decisions depend on utilities, which may include intrinsic biomechanical, oculomotor, as well as cognitive costs. To measure these utilities, we introduce a new saccadic preference elicitation experiment in which subjects repeatedly selected one out of two alternative gaze targets. This allowed measuring subjects’ individual saccadic preferences independently of spatial image properties in terms of three factors: saccade amplitude, direction of saccade relative to the preceding gaze shift, and global gaze direction. Importantly, subjects were instructed to freely select one of the two alternative targets. This elicits their intrinsic decision making process and thereby allows observing their preferences, which can be used to infer the underlying subjective utilities.

While the notion of a utility function (Von Neumann & Morgenstern, 1944) and its estimation from a set of binary decisions has been extensively studied in economics, psychology and machine learning (Friedman & Savage, 1952; Thurstone, 1927a, 1927b), we adopt a class of models called random utility model (RUM) (McFadden et al., 1973; Train, 2009), which allows explicitly modeling the uncertainty of subjective utilities. Specifically, we devised several RUMs differing both in the factors contributing to the decision as well as in the type of uncertainty about utilities. While popular RUMs involve normally distributed uncertainties about utilities, we derived a RUM with log-normally distributed uncertainty, allowing for Weber-Fechner type phenomena. In the spirit of rational analysis (J. R. Anderson, 1991; Gershman et al., 2015; Simon, 1955), we subsequently invert these utility models of human saccadic behavior through Bayesian inference of parameters describing individual subjects. Finally, we perform model comparison using the Bayesian Information Criterion (BIC) and find overwhelming evidence in favor of a RUM involving all three saccadic features and log-normally distributed uncertainty.

The results show, first, that subjects shared similar preferences for shorter amplitude saccades in line with previous research in motor control. Second, all participants shared a preference for saccades in the direction back to the previous gaze location, contrary to numerous reports on “inhibition of return” (Bennett & Pratt, 2001; Klein & MacInnes, 1999). Surprisingly, the experiment revealed marked differences in individual preferences for global gaze directions, which were stable across most participants when measured in a second experimental session after one week. Taken together, we provide the first computational account of individual differences in saccadic preferences based on a neuroeconomics approach allowing to identify factors contributing to the utility of saccadic gaze movements independent of complex image content and to dissociate factors shared across subjects from those underlying individual variation. Our results are not only important to understand the intrinsic contributions to saccadic choices but are also of broad consequence to computational models of gaze selection. Finally, both the experimental code as well as the estimated utility functions are released as part of this publication, allowing the research community to factor in the cost of a gaze shift in experiments involving saccadic eye movements.

## 2 Results

### Saccadic preference elicitation experiment

A total of N=14 subjects took part in the experiment and completed on average a total of approximately 5000 saccadic decisions each. Participants sat in front of a monitor approximately subtending an angle of 52° horizontally and 31° vertically. Each trial started with the appearance of a central fixation target, which subjects had been instructed to look at. After fixating the central target for a delay of random duration between 0 and 0.5 seconds two saccadic targets appeared at random locations on the screen and subjects chose freely to saccade to one of them, see Fig. 1a for a schematic. After choosing one of the two targets by carrying out a saccadic eye movement to the preferred target, the central fixation target reappeared whereby the subsequent trial started, as illustrated in Fig. 1b. Subjects repeated the experiment in a second experimental session, which was scheduled on average 6.4 days after the first session.

**Figure 1.**
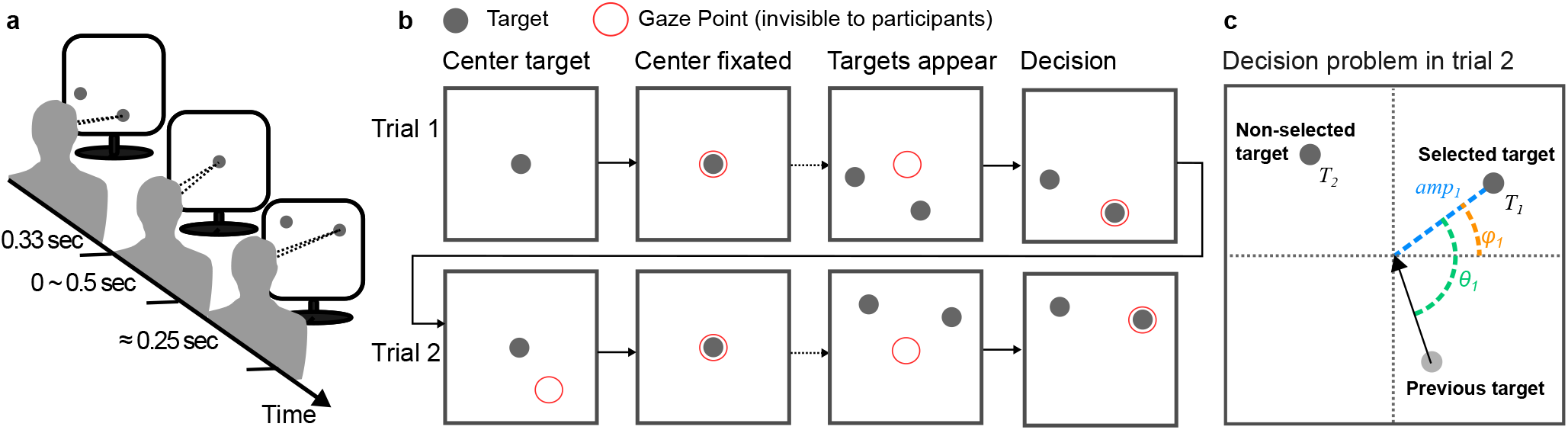
Saccadic preference elicitation task. **a** Experimental setup, with the subject in front of a monitor, selecting targets, with his/her gaze. **b** Illustration of experimental sequence for two consecutive trials. Each trial starts with appearance of the central fixation target. After the subject fixates the target, two alternative targets appear after a variable delay between 0 and 0.5 sec. The subject then decides for one of the two available targets by fixating. This terminates the trial and initiates the subsequent trial. **c** One 2AFC decision and visualization of the features determining the decision process. The arrow indicates the last saccade. For visual clarity features are only shown for target 1.

### Saccadic choice features

To quantify the saccadic preferences of subjects, we transformed the raw positions of each of the two simultaneously displayed targets from screen coordinates to three features, which in the past have been argued to influence eye movement decisions. The first feature is the amplitude *amp* of the potential saccade, measured as visual angle from the central fixation target. The second feature is the global direction *ϕ* of the target within the visual field, calculated as the counterclockwise angle between the horizontal, positive x-axis and the target’s position. Thus, an angle close to 0° corresponds to targets on the horizontal direction to the right and an angle of 90° corresponds to a target up on the vertical direction. The third and final feature is the change in direction *θ* relative the previous gaze shift. This can be defined as the counterclockwise angle between the position of the last chosen target and the position of the potential next target of the upcoming saccade. Accordingly, a relative angle of 0° (or 360°) corresponds to a saccade back in the direction toward the previous saccadic target, a so called return saccade, while a relative angle of 180° corresponds to a target in the opposite direction of the last gaze target, i.e. a continuation in the same direction as the previous gaze shift, a so called forward saccade. These three features are visualized for one target in an example trial in Fig. 1c.

### Behavioral Results

To analyze how the above defined three features influenced participants’ saccadic decisions, we computed the proportion of targets chosen at equally spaced intervals of each feature. This choice allows to express piecewise linear dependencies, which balances expressibility of functional relationships without assuming specific functional forms. The resulting proportions are shown in Fig. 2a-c. First, the amplitude of a potential saccade had a clear influence on the proportion of chosen targets with an approximately linear increase in radial distances leading to a linear decrease in proportion of chosen targets, see Fig. 2a. The linear regression between the radial distance and the average proportions of chosen targets was highly significant with *β* = −0.0611; 95% CI = (−0.0634, −0.0588); t(9) = −59.53; *p* < 0.001. The corresponding *R*^2^ value was 0.9975.

**Figure 2.**
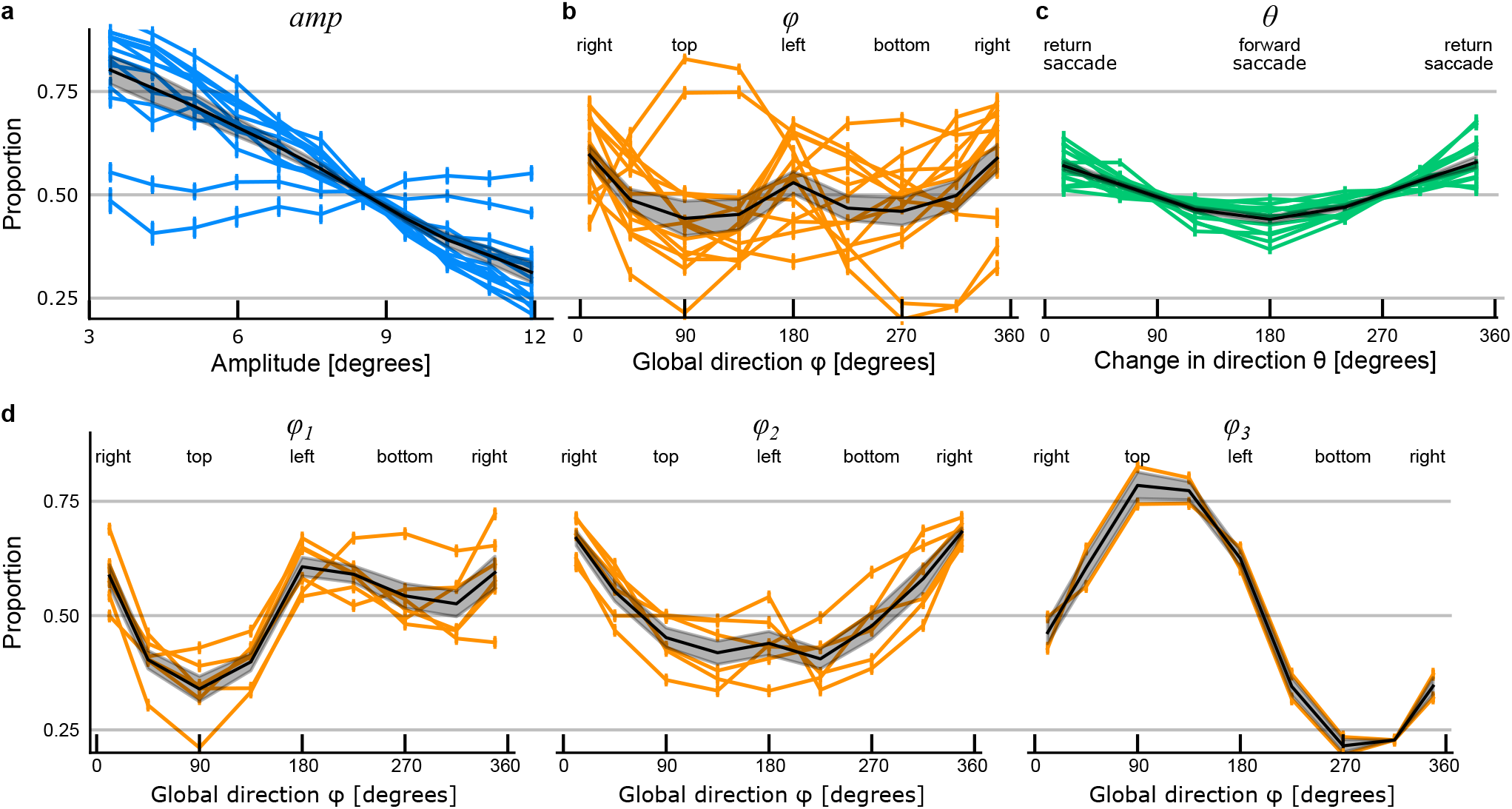
Behavioral results. **a-c** The proportion of chosen targets for the three saccadic features amplitude (**a**), direction (**b**), and change in direction (**c**). Colored curves represent single subjects’ data, while the black curves show the mean over all subjects. The error bars for the colored curves and the shaded area for the black curve indicate ± one standard error of the mean. **d** Proportion of targets chosen for different directions (as in subplot **b**), grouped by the cluster found by k-means clustering with *k* = 3. For all of the 3 plots, every curve represents one subject.

The proportion of saccadic choices was also consistently influenced by the change in direction of a gaze shift in that all subjects preferred targets located in a similar direction as the previous gaze target, i.e. targets with *θ* values close to 0°or 360°. Accordingly, targets located opposite in direction to the previous gaze location, i.e. continuing in the same direction as the previous gaze shift, were least preferred with an approximately linear decrease in choice probability with an increase of the relative angle towards 180°, see Fig. 2c. The strength of this effect varied between subjects but was highly significant: *β* = 0.0009; 95% CI = (0.0007, 0.0011); t(6) = 11.9783; *p* < 0.001. Thus, the amplitude and change in direction of saccadic targets resulted in similar effects on the proportion of targets chosen for most subjects, albeit with differences in their strength across participants.

The average influence of the global direction on the probability of target selection across subjects showed a significant preference for horizontal saccades (t(141284) = 21.7435; *p* < 0.001), consistent with previous reports. But, this influence showed considerable variation across subjects. The proportions of targets chosen across different global directions for all subjects are shown in Fig. 2b. Because of the large idiosyncratic differences, there are no significant differences between targets at the right (≤45° or >315°), the top (>45° and ≤135°), the left (>135° and ≤225°) or at the bottom (>225°and ≤ 315°). Differences between the proportions chosen at those respective locations are not significant at the *α* = 0.01 level (ANOVA: F(3,52) = 2.9448; *p* = 0.0414). To analyze the different groups, k-means clustering with different numbers k clusters were computed. Using the elbow-method of the within cluster sum of squared errors (Thorndike, 1953), the optimal number of clusters was determined to be k=3, (sum of squares was (*k* = 1 : 1.8452, *k* = 2 : 0.7480, *k* = 3 : 0.3921, *k* = 4 : 0.2591)). The three clusters were consistent over 10 runs with different random initialization values. The proportions of chosen targets for the three clusters are shown in Fig. 2d. The three groups of subjects can be described as having an aversion for the top half of the display (left plot), a preference for the right side of the display (middle plot), or for the top half of the display (right plot). We repeated the ANOVA described above for each of the three groups obtained through clustering to investigate the difference in preferences between cardinal directions, which was highly significant: (*left*: F(3,22) = 18.2074; *p*< 0.001, *middle*: F(3,22) = 17.6914; *p*< 0.001, *right*: F(3,5) = 86.6257; *p* < 0.001).

The strength of the three factors amplitude, global direction, and change in direction on saccadic preferences can be quantified in terms of the difference between the minimum and the maximum proportion of target selection. The target’s amplitude has the strongest influence on the difference in proportion of chosen targets (mean=0.5046, SD=0.1827), followed by the global direction (mean=0.3627, SD=0.1087) and the change in direction with the smallest influence on choices (mean=0.1289, SD=0.0664). The relative strength of the influence of these three factors varies moderately between subjects, particularly the influence of global directional preference had a maximal value of 0.6300 (max=0.8258, min=0.1958) and a minimal value of 0.2352 (max=0.6647, min=0.4296). This variability is illustrated in Fig. 2b and d.

### Stability of preferences across time

All previous analyses aggregated data of each participant from both experimental sessions. We furthermore investigated, how the three target features amplitude, global direction, and change in direction influenced the proportion of chosen targets across the two experimental sessions, which were on average 6.4 days apart. We observed overall very small differences between the preferences of both experiment dates for all participants. To quantify the stability of preferences, we calculated a MANOVA between the first and the second experiment date. The MANOVA for all three features were not significantly different at the *α* = 0.1 level: (**amp**: F(11,16) = 1.7199, *p* =0.1573; *ϕ*: F(9,18) = 0.6050, *p* = 0.7771; *θ*: F(7,20) = 1.2728, *p* = 0.3127).

### Preferences and decision time

The experimental design involved a variable temporal delay sampled uniformly at random between zero and half a second between the onset of fixation at the central fixation target and the appearance of the two saccadic targets (the time between the second and the third column in Fig. 1b). This was implemented to allow examining the time subjects spent fixating the central target before initiating the subsequent saccade and investigating the relationship between saccadic preferences and saccadic timing. Accordingly, this temporal delay includes the time required for the decision-making process of which saccadic target to choose and it can be used to compare the proportion of decisions depending on the decision time. Subjects’ decision times were approximately constant for all central fixation durations longer than 250 milliseconds (mean=246.7709, SD=1.5280, ANOVA: F(15,36584) = 0.5493; *p* = 0.9138), see also Supplementary Fig. S3. For shorter delays, the time before initiating the saccade increased approximately linearly up to 285 milliseconds (*β=* −0.1525; 95% CI = (−0.1713,-0.1337); t(12) = −17.5237; *p* < 0.001). Therefore, we divided decisions based on their central fixation duration into short (<250ms) and long (≥250ms). We repeated the analysis of the stability of saccadic preferences in terms of the three features for short and long decision times. Again, we calculated a MANOVA for each feature and all three analyses were not significantly different, at the *α* = 0.01 level: (**amp**: F(11,16) = 0.8375, *p* = 0.6094; *ϕ*: F(9,18) = 0.1171, *p* = 0.9988; *θ*· F(7,20) = 3.6045, *p* = 0.0113). Thus, we do not find evidence for a difference in the intrinsic preferences depending on the duration of the saccadic decision time.

### Random utility model of saccadic preference

While the above analyses empirically quantify the likelihood of subjects’ choices, they do not provide a computational account of saccadic decisions. Therefore, we devised a computational model of saccadic choices by conceptualizing saccadic gaze shifts as a decision process (Gottlieb, 2012; Hayhoe & Ballard, 2014; Hoppe & Rothkopf, 2019). Specifically, we devise a set of saccadic RUMs starting from classic RUMs (McFadden et al., 1973; Train, 2009), which assume that individual choices are determined by the comparison of latent utilities of alternatives. In case of a decision between two targets *T*_1_ and *T*_2_, the respective latent utilities *U*(*T*_1_) and *U*(*T*_2_) are compared internally by computing their difference *U_d_* = *U*(*T*_1_) – *U*(*T*_2_). The decision maker makes decision *y* by either selecting target *T*_1_ in case this difference is positive or target *T*_2_ otherwise. However, RUMs assume that utilities are noisy and therefore the decision is based on a noisy difference of utilities *z* = (*U*(*T*_1_) + *δ*_1_) – (*U*(*T*_2_) + *δ*_2_) = *U_d_* + *ϵ*, where *z* is the decision variable and *ϵ* can be though of as a noise term rendering the decision stochastic resulting from the individual noise values *δ_i_*. Thus, the decision is based on evaluating whether the decision variable *z* is larger than zero, which is customarily expressed with the so called indicator function 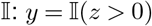.

If the uncertainty underlying *δ_i_* for each target is modeled as being normally distributed, then the difference in utilities *U_d_* is also normally distributed, leading to the classic probit model (McFadden et al., 1973; Murphy, 2021; Train, 2009). In this case, the probability *P*(*y* = 1) of the decision maker selecting as its choice *y* the target *T*_1_ is simply the expectation of the indicator function over all possible values the decision variable can attain. This can be expressed with the following integral, which leads directly to an expression involving the cumulative normal distribution Φ(·):

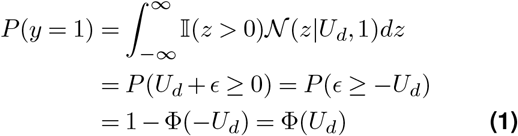

The functional form of the utility values and possible alternative probability distributions modeling the utilities’ uncertainty depend on the respective decision problem. In the following paragraph, we explain our respective choices. Regarding the utility function, we made two assumptions. (1) The utility of a target does only depend on the amplitude, the direction and the change in direction of the possible saccade and (2) the influence of every feature can be described independently of the other ones. The second assumption is known as *additive utility independence*. Accordingly, the utility function *U*(*T*) for a saccadic target *T* is defined as the sum of three independent functions, each applied to one of the three features:

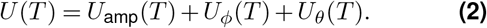

Based on the empirical results described in the previous sections, we chose *U*_amp_ to be linear in the amplitude, *U_θ_* to be linear in the deviation from an angle of 180° and *U_ϕ_* to be a piecewise linear function with 6 equispaced fixed turning points. Additional details for all three functions and their design decision can be found in the Methods section. As for the shape of the uncertainty, the classical probit model assumes normal randomness in the latent utilities as a way to capture factors influencing the decisions, which are unobserved to the researcher. Here, we additionally derived a RUM under the assumption that utilities may be internally uncertain. Accordingly, we use log-normally distributed uncertainty in the utility of each target, since this choice for the internal representational uncertainty can accommodate Weber-Fechner phenomena, which are pervasive in perception and action. While it is known that there is no closed form for the distribution of *ϵ*, which in this case is a difference of two log-normally distributed random variables, we utilize numerical approximations to compute its value. Further details on this numerical approximation are provided in the supplementary material. With this approximation, we now have a way of calculating the probability for a target to be chosen given the respective utility values and under the assumption of independent trials. A schematic illustration of the RUM, as well as the stochastic decision process is provided in Figure 3a. Further details on how to invert these models and perform Bayesian inference over the models’ parameters involving probabilistic programming is given in the Methods section.

**Figure 3.**
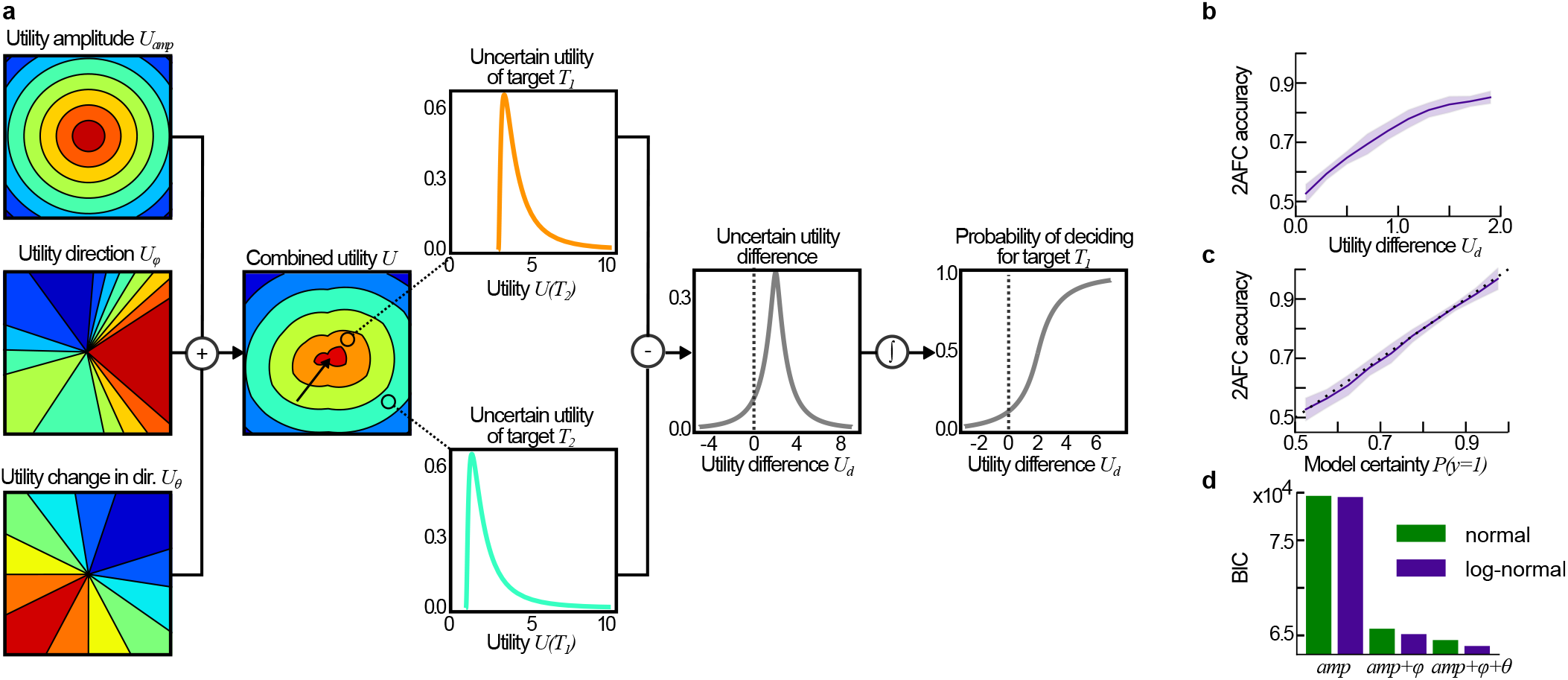
Saccadic random utility model. **a** The RUM can generate global preference maps for each feature individually, which can be combined by summation. A binary decision can then be modeled by taking the respective utility values at the target locations and adding the respective uncertainty. The integral of the uncertain utility difference up to 0, i.e. the area under the curve up to the dotted line in the right most plot, is the probability for choosing target *T*_1_. **b** The model’s accuracy as function of the difference in utility between targets, and **c** the probability assigned to the chosen target. In both plots, the mean over all subjects is indicated by purple curve, while ±1 standard deviation over all subjects is shown as the shaded area. **d** BIC scores for the six RUMs. The models included either only the utility of the amplitude *amp* of an upcoming saccade, the amplitude *amp* and the global direction *ϕ*, or the amplitude *amp* together with both the global direction *ϕ* and the change in direction *θ*. These three factors were paired with normal uncertainty about utilities and log-normal uncertainties, respectively.

### Model comparison and performance

To investigate, whether all saccadic features are relevant for describing participants’ choices, we calculated the Bayesian information criterion BIC (Schwarz, 1978) for RUMs involving (1) only the amplitude, (2) the amplitude and the direction and (3) all three features. In addition, we also investigated whether the behavioral data is better accounted for by utilities with normal or log-normal uncertainty. With these two variations, we have 6 models in total. For each model, we calculated the respective BIC score by evaluating each model on the data of all subjects. The resulting BIC scores are depicted in Figure 3d. The difference between models with normal and log-normal uncertainty distributions is consistently in favor for the model with log-normal subjective preference uncertainties. For both uncertainty distributions, each additional feature decreased the BIC score, proving evidence that the performance of the model depends on all three features. The best fitting model, the RUM with all three features and log-normally distributed uncertainty, had a difference in BIC score to the second best fitting RUM of ΔBIC=621.09, which can be considered overwhelming evidence. To evaluate the accuracy of saccadic predictions of the RUM, we compared the participants’ saccadic decisions with the predictions of the model. Model choices were calculated by always deciding for the target with the higher utility. For all participants the accuracy is approximately 80% (mean: 0.7915, SD: 0.0502, min: 0.6889, max: 0.8615), i.e. 56203 of a total of 70643 saccades in the experiment were predicted correctly by the RUM. Figure 3b and c extend this analysis, by showing the accuracy of the model’s predictions dependent on the utility difference between both saccadic targets (b) and the model’s probability of being correct (c). The latter probability was calculated using the approximated cumulative distribution of the difference of two log-normal distributions, i.e. as Ψ(|*U_d_*|). These two plots show how we can convert utilities to decision probabilities and how well the model is calibrated.

### Individual saccadic utilities

Parameters describing saccadic utilities on the basis of the RUM with log-normally distributed uncertainties were obtained for each individual subject. Figure 4a shows a summary of the inferred parameters. On average, the utility weight for the amplitude has the largest absolute value, reflecting the fact that the amplitude has the strongest influence on saccadic decisions. Since larger saccade amplitudes are less preferred, the value of *w*_amp_ is negative. The parameters describing the influence of the saccadic direction *w_ϕ_* show a wide spread, agreeing with the empirical observation of idiosyncratic differences across subjects both in the directional preferences and the strengths of preferences. Finally, the parameter values of *w_θ_*, quantifying the influence of the change in saccadic direction, have positive values close to 0, also in line with the empirical observation of the preference for return saccades from Figure 2c, which is weaker than the preferences for the other two features. Overall, the RUM results confirm the empirical observations including the order of strength of the influence of the three saccadic target features shown in Figure 2.

**Figure 4.**
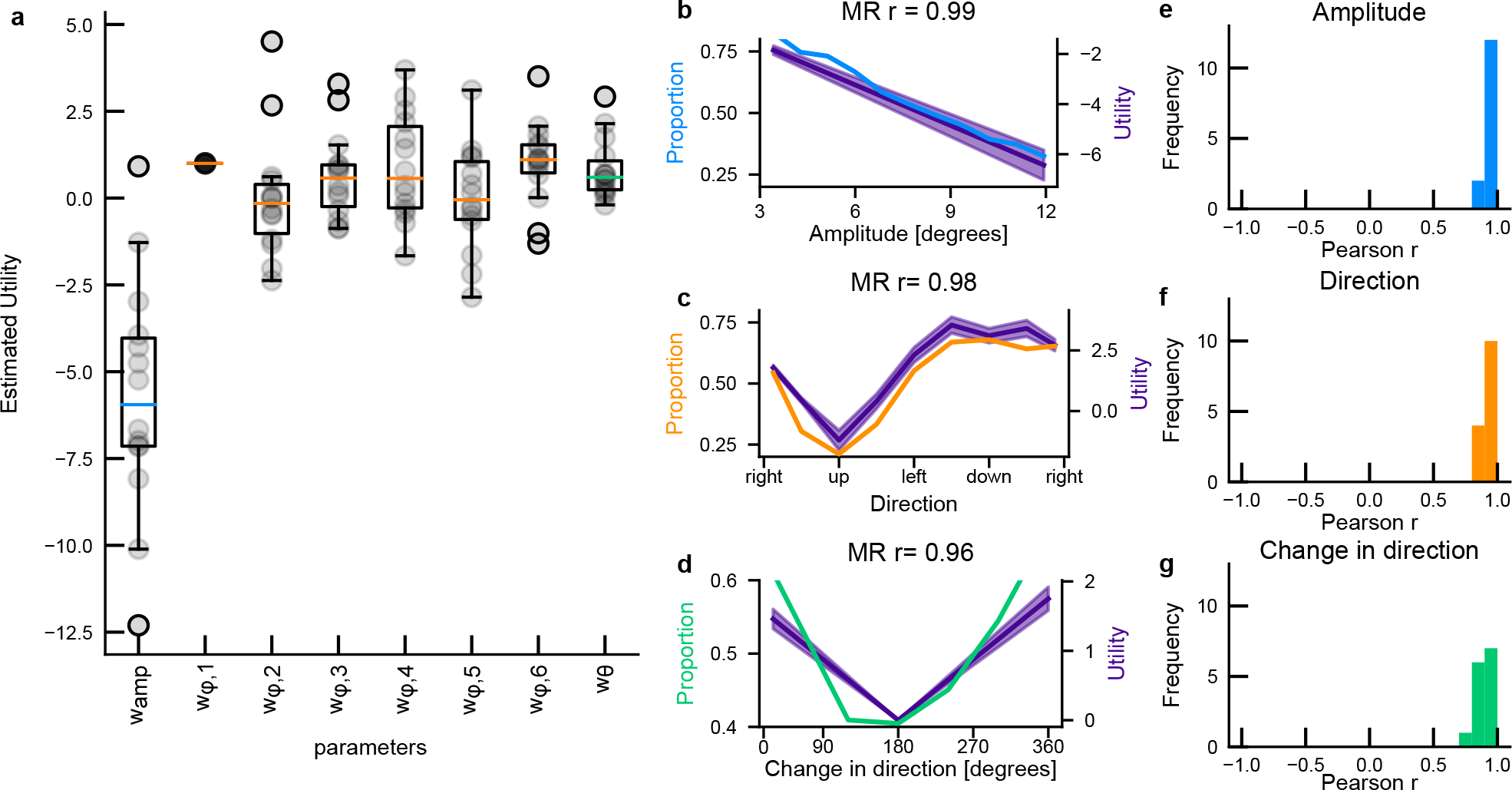
Modeling results. **a** Boxplots for parameter values, showing the distribution of those over all subjects. The orange lines indicate the median of the respective parameter. *w*_*ϕ*,1_ is required to calculate the utility, yet it would be redundant to estimate it, thus it was fixed to a value of 1 for all subjects. Details regarding all parameters are in the Methods section. **b-d** Comparison model utility values with subjects’ preferences, for amplitude (**b**), direction (**c**) and change in direction (**d**). Each plot shows the comparison between subjective preferences for one specific feature and the respective utility values. The title indicates the subject and the Pearson correlation coefficient between the two lines. The shaded area around the purple line visualize the mean ± 2 standard deviations of all parameters. **e-g** The histogram of Pearson correlation coefficients over all subjects, between utility values and preferences for amplitude (**e**), direction (**f**) and change in direction (**g**).

Examples of how the predicted utility values compare to the respective empirical proportions for one subject is shown in Figure 4b-d, which includes the uncertainty in posterior inferences about the parameter estimates obtained with Bayesian inference. To this end, model utility values were calculated for each preference interval. Note that utility values for single features, independent of the others can only be calculated because our utility function was constructed with the additive utility independence property. The plots give a first indication that the RUM has low uncertainty about the parameters best describing subjects’ preferences, a fact that can be attributed to the large number of decisions compared to the number of parameters for each subject. To compare whether utility values and proportions match, we calculated the Pearson correlation coefficients. For the specific subject shown, all coefficients are larger than 0.95. Figure 4e-g then validates that the match between subjective preferences and model utility values is indeed very good for all subjects and features. The correlation coefficients for all features and subjects are above 0.7 (min=0.7256, max=0.9982, median=0.9733, mean=0.9383), showing that the RUM not only predicts the correct targets well but also the respective preferences for the three features. It is important to note that the subjective preferences could in principle display more complex behavior than the chosen parameterization of utility functions, because the empirical functions we used are more expressive compared to the parameters entering the RUM. Yet, the fact that modeled utility values and subjective preferences do align well verifies the modeling assumptions.

## 3 Discussion

Previous research on human active gaze selection has reported that not only certain locations within a scene are more likely to be looked at (Borji et al., 2013; Findlay & Gilchrist, 2003; Hayhoe & Ballard, 2005; Henderson, 2003; Navalpakkam et al., 2010; Rothkopf et al., 2007; Schütz et al., 2012; Torralba et al., 2006; Yarbus, 1967) but that also certain gazeshifts are more likely than others. The problem with the approach of empirically estimating both these likelihoods from human gaze recorded while looking at images is that they are not independent of each other as demonstrated by simply rotating the images (Foulsham et al., 2008). And indeed, investigations of oculomotor biases have reported a multitude of puzzling and in part contradictory observations (A. J. Anderson et al., 2008; Bays & Husain, 2012; Bennett & Pratt, 2001; Dodd et al., 2009; Gilchrist & Harvey, 2000; Horowitz & Wolfe, 1998; Klein & MacInnes, 1999; Luke et al., 2014; Peterson et al., 2001; Smith & Henderson, 2009; Tatler & Vincent, 2009; Wilming et al., 2013). Here, we conceptualized gaze shifts as a sequential decision-making process (Gottlieb, 2012; Hayhoe & Ballard, 2014; Hoppe & Rothkopf, 2019), which has the consequence that intrinsic behavioral biases in the selection of visual targets can be quantified in terms of subjective utilities. This approach is motivated by the abundant literature on the tight connection between eye movements and decision making in the brain (Gold & Shadlen, 2007; Gottlieb, 2012; Hayhoe & Ballard, 2005), motor control relating gaze shifts to movement costs (Harris & Wolpert, 1998; Todorov & Jordan, 2002), and the recent literature on modeling behavioral and cognitive costs in eye movements (Hoppe & Rothkopf, 2016, 2019; Lisi et al., 2019; Petitet et al., 2021). In this framework, individual gaze shifts integrate behavioral benefits of prioritizing certain locations of the visual scene with the intrinsic costs of carrying out certain eye movements. This allows adopting principled methods from neuroeconomics (Glimcher & Fehr, 2013; Körding et al., 2004), to measure the subjective utilities of saccadic gaze shifts. Accordingly, we devised a new saccadic preference elicitation experiment in which we measured participants’ saccadic choices independently of complex spatial image statistics in terms of three features: the gaze shift’s amplitude, its direction within the visual field, and its change in direction relative to the previous saccade.

Along with the new saccadic preference elicitation experiment, we devised RUMs (McFadden et al., 1973; Murphy, 2021; Train, 2009) quantifying individual participants’ subjective utilities of a gaze shift. The models assume that uncertain internal representations of the utilities of alternative gaze targets are compared and the decision is made for the target with higher utility. All models assume additive utility independence for the three contributing factors and differed in both the included features contributing to saccadic utility and the form of the utilities’ uncertainty. Model parameters were estimated for each individual subject and model comparison through the BIC showed that subjects’ saccadic decisions were best described by a RUM model involving all three features and log-normal internal uncertainty. In summary, approximately 80% of all gaze shifts across subjects are predicted by the best fitting RUM. Importantly, the RUM allows quantifying the relative contributions of the three features to individual saccadic decisions and it allows dissociating factors that are shared across subjects from those that underlay idiosyncratic variation. In the following sections we discuss these three factors individually.

All subject showed a consistent linear influence of saccadic amplitude on utility, albeit to varying degrees. This result agrees with previous empirical studies, which have reported that subjects appear to prefer shorter to longer saccades in a variety of tasks, a property that has been referred to as “proximity preference” (Engel, 1971; Koch & Ullman, 1987). As an example, an analysis of a large collection of gaze shifts showed that the proportion of saccades decreases linearly with distance (Tatler et al., 2006). This linear dependence may be explained at the implementational level based on the well known signal dependent variability of neuronal firing. Models of optimal motor control (Harris & Wolpert, 1998; Todorov & Jordan, 2002) have used criteria such as the minimization of saccadic endpoint variance after an eye movement under the influence of signal dependent neuronal noise to plan optimal saccades. These studies conclude that larger saccades go hand in hand with a linear increase in endpoint variability, thereby leading to a preference for shorter saccades.

All subjects in our experiment shared a consistent preference for eye movements directed back to the previous location of gaze, so called return saccades. This preference was gradual such that the subjective utility of a saccade decreased approximately linearly with an increase of angular deviation from the direction to the previous gaze target. This component of participants’ utility was the most consistent of the three factors we investigated, both across subjects and across the two experimental repetitions, albeit smallest in magnitude. This result may seem particularly surprising, given numerous reports on the likelihood of return saccades being lower than expected by chance. This so called phenomenon of spatial inhibition of return (Bennett & Pratt, 2001; Klein & MacInnes, 1999), particularly prominent in studies involving naturalistic images (Bays & Husain, 2012; Luke et al., 2013; Luke et al., 2014; Wilming et al., 2013), has been interpreted as an oculomotor bias encouraging active exploration of visual scenes or as “foraging facilitator” (Klein & MacInnes, 1999; Wilming et al., 2013). In the present experiments, subjects neither carried out a search task, nor did the visual scene contain complex, natural scenes. Accordingly, the phenomenon of spatial inhibition of return cannot be attributed to an intrinsic oculomotor bias. Instead, the saccadic system favors decisions that result in smaller rather then larger change in direction relative to the previous gaze shift. Thus, any observed deviations from this pattern needs to be attributed to a separate process, consistent e.g. with the notion that only certain tasks elicit a spatial inhibition of return behavior (Dodd et al., 2009).

At first look, the preference for the global direction of saccades averaged across subjects showed the often reported pattern of a significant bias favoring horizontal gaze shifts (Foulsham et al., 2008; Gilchrist & Harvey, 2006), albeit to a lesser degree. However, the cause for this bias has been debated controversially and previously reported results (Foulsham et al., 2008) clearly demonstrated, that simply rotating the image leads to a change in the distribution of preferred saccade directions. Thus, the observed higher likelihood of horizontal saccades cannot solely be attributed to oculomotor biases, as noted previously (Tatler & Vincent, 2009). In the current experiment, the targets were distributed equiprobably across all orientations and complex image statistics were not present. Thus, the observed biases can be attributed to participants’ intrinsic directional preferences. However, the directional preference obtained as average over individual participants hides the significant idiosyncratic differences in global gaze direction preference across subjects. Clustering these preferences revealed idiosynchratic interindividual differences, which can be attributed to three different preference groups. Previous research has observed marked individual differences in oculomotor behavior (Bargary et al., 2017; Henderson & Luke, 2014; Risko et al., 2012) and recently individual differences in the selection of semantic image content (De Haas et al., 2019) have been reported. Our results show that such individual differences extend to the preference in saccadic gaze shifts for global directions within the visual scene.

The inferred preferences for all three saccadic features estimated from the behavioral data were remarkably stable for most participants across a week as observed in a second experimental session. Pearson correlation coefficient of preferences were consistently higher within subject across the two experiments than across subjects. To exclude the possibility, that these preferences were somehow learned during the experiment, we investigated, whether subjects’ preferences changed during the early stages of the experiment. To test this hypothesis, we compared the preferences estimated from the entire experimental data (see Figure 2) with the data from the first 10 blocks of the experiment. A MANOVA between the first 10 and the all blocks, resulted in no significant differences at the *α* = 0.05 level (**amp**: F(11,16) = 0.8383, *p* = 0.6088; ***ϕ***: F(9,18) = 2.0474, *p* = 0.0936; ***θ***: F(7,20) = 0.9150, *p* = 0.5153). For additional comparisons, see also Figure S4. Including only 10 minutes of data from each subject already shows remarkable similarities to the overall preferences, providing evidence that the reported results are indeed attributable to internal saccadic decision preferences, instead of being learned.

Beyond quantifying the spatial preferences of saccadic decisions, the current experimental design additionally allowed investigating the dependence of these decisions on the time available to subjects before carrying out the eye movement. Previous research has reported that the duration of a fixation before a return saccade to the previous fixation location is prolonged compared to a saccade continuing in the same direction, the so called “temporal inhibition of return” (Hooge et al., 2005; Klein & MacInnes, 1999). Further studies found that the fixation duration was longer the larger the angle of the subsequent saccade was relative to the direction of the forward saccade, i.e. the larger the deviation from continuing in the same direction as the previous gaze shift (A. J. Anderson et al., 2008; Smith & Henderson, 2009; Wilming et al., 2013), the so called “saccadic momentum”. In the present experiments, we did not find a significant influence of the fixation duration on the preferences of the subsequent saccade for any of the three saccadic features amplitude, gaze direction, and change in gaze direction. Thus, temporal inhibition of return is not an intrinsic oculomotor bias but may be elicited under particular stimulus and task conditions.

The RUM model provides fundamental advantages in quantifying saccadic decisions. First, instead of reporting oculomotor biases in terms of empirical likelihoods of certain saccadic parameters, the RUM quantifies preferences for saccadic decisions in terms of utilities for saccadic features. This allows framing saccadic decisions with a rigorous computational framework of decision making and neuroeconomics. Second, individual utility values allow quantifying trade-offs between different features. As an example, participant *NV* subjectively equates a direction 60° shift, from bottom to bottom right, with shifting the saccade approximately 90° towards a return saccade, while participant *BS* values a shift from downward to upward saccades equally as an increase in 3.5°amplitude. Third, the utilities characterized in the present study allow not only estimating the likelihood of individual gaze shifts but also their relative probability, as the RUM with log-normal uncertainties is well calibrated. Fourth, the RUM allows quantifying the utilities of all potential gaze targets within the visual field of view in terms of a saccadic utility map, see Figure 3. Fifth, the utilities found in this experiment can be used to factor in the cost of a gaze shift in experiments involving saccadic eye movements, which have often treated all gaze shifts as being equally likely. As an example, experiments on visual search (Gilchrist & Harvey, 2000, 2006; Horowitz & Wolfe, 1998; Peterson et al., 2001; Torralba et al., 2006) often assume, that saccadic gaze shifts are essentially for free and equiprobable in all directions. Finally, while computational models of human gaze selection such as “saliency models” have made extensive use of a generic process of “inhibition of return” (Borji & Itti, 2012), the current results suggest that a more accurate prediction of human gaze may be obtained by employing the actual human preferences for saccadic eye movements.

In conclusion, we introduced a saccadic preference elicitation experiment in the framework of neuroeconomics, which made it possible to reason about the influence of saccadic features on their probability to be made in terms of intrinsic utilities. The saccadic RUM captures the effect of amplitude, direction and change in direction on the decisions of gaze shift. We found common linear preferences for shorter and return saccades and idiosyncratic preferences for the direction. Finally, we formulated a RUM, which proved to be able to predict those preferences and makes it possible to include the results of this study into experiments involving gaze selection and gaze prediction models. On a more conceptual level, the current results further show the importance of taking the intrinsic costs and benefits of behavior into account, even in behaviors seemingly only conceptualized as perceptual acts.

## 4 Methods

### Participants

14 subjects (7 females and 7 males) took part in the experiment in exchange for course credit or monetary compensation. Participants’ age ranged from 19 to 33, mean=23.93, median=22.5. All subjects signed a consent form prior to the experiment. The experimental procedure was approved by the Ethics Committee of the Technical University of Darmstadt and was in accordance with the declaration of Helsinki. All subjects had normal vision in order to ensure good trackability of the eyes during the experiment.

### Task design and procedure

Participants were continuously presented with repeating two-alternative forced-choice (2AFC) trials. The two alternative targets were displayed after a delay chosen uniformly at random between 0 and 0.5 seconds and subjects indicated their choice by saccading to the preferred target. This choice was followed by a delay of one third of a second. After each decision, subjects had to fixate the newly appearing target at the center of the screen, thereby initiating the subsequent trial. Two consecutive trials are illustrated in Figure 1b. The following specifications in visual degree are based on an average viewing distance of 55 centimeters of our seated participants. Small deviations in viewing distance dependent on individual subjects and time over the experiment were unavoidable. Target locations were sampled uniformly at random from a circle around the center of the screen with a radius of 12 degree (412 pixels). Targets closer than 2.9 degree (100 pixels) were resampled. Overall, each participant completed two sessions of the experiment, which were at least four days apart (mean=6.4 days, SD=2.4, min=4, max=14). In each session, subjects performed 45 blocks of 1 minute length. During each block, subjects were instructed to complete as many trials as they could. After each block, subjects were provided with feedback about the number of saccadic decision performed during the last ten blocks and were free to either take a short break or start the next block by pressing a key on a regular keyboard. A five minute break to prevent fatigue after 25 trials was mandatory.

### Stimuli and apparatus

Targets were circles with a smaller concentric circle. The outer diameter was 0.7 degree, the inner was 0.1 degree, and the target had a mean luminescence of 3.4*cd/m*^2^. The background was dark gray and had a mean luminescence of 33.2*cd/m*^2^. A target was counted as fixated if the eyetracker detected a gaze location closer than 1.4 degree (50 pixels) to the center of the target. If a fixation was within this distance to both targets, only the closer target was regarded as being fixated. The fixation duration necessary to collect the central fixation targets was drawn uniformly at random between 0 and 1/2 of a second for each trial. For any 2AFC target, this time was constant at 1/3 of a second. The experiment was presented on a *DELL P2314H* monitor with a diagonal of 58,42cm (23”), a resolution of 1920×1080 pixels, and 60Hz refresh rate. Gaze behavior was tracked with a *SMI Red* eyetracker with a sampling frequency of 250 Hz. Participants were calibrated using 5 point calibration. The calibration was repeated if the average tracking error was above 1.0 degree or the tracking error on a single point was above 2.0 degree. For overall better trackability of the gaze, subjects’ placed their heads on a chin rest.

### Data processing

Equations 3–5 show the actual formulas with which the three features described above were calculated from raw screen coordinates:

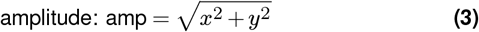

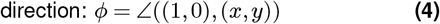

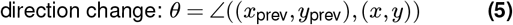

The data stream from the experiment was grouped by trials, where each trial started when the central target appeared. Possibly unfinished trials at the end of each block were excluded, as well as the first trial of each block, because it has no preceding gaze shift. All remaining data points were treated as a single decision with their respective features.

Subsequent analyses of the empirical proportions of gaze selection were carried out by binning decisions by their respective features. All analyses and corresponding plots involving interval features use the same binning parameters. For the amplitude, we used twelve equally spaced intervals between 2.9° and 12° of visual angle (100 pixels and 412 pixels). The edges of the direction intervals were: 0°,22.5°,67.5°,112.5°,157.5°, …, 360°, because this ensures the center of the interval to be the exact horizontal, vertical and diagonal directions. The change in direction angles were divided into six equally spaced intervals of 60° width each.

### Random Utility Models

The statistical modeling was done with a random utility models (RUM) (McFadden et al., 1973; Murphy, 2021; Train, 2009). Classic RUMs assume that the decision maker is influenced by observable factors and factors, which are unobserved by the experimenter. Thus, the stochasticity in decisions is the result of unobserved factors, which are modeled as jointly introducing normally distributed noise(McFadden et al., 1973; Train, 2009). More generally though, the stochasticity in decisions may also be attributed to uncertainty(Murphy, 2021), which may include internal representational sources. Different from common RUMs, we derived a new model with the log-normal distribution for the internal noise on the value of each target, accommodating Weber-Fechner type variability. Specifically, each error term was distributed with 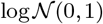. As common in RUMs, the choice of a standard deviation of 1 is arbitrary. Intuitively, the probability of choosing a specific target does only depend on the fraction of the noise parameter and the utility difference. This means, that a model that doubles its noise parameter and all utility differences, by just doubling all linear weights, is exactly the same model as the one without doubling all parameters. Hence, it is common to fix the variance parameter of the distribution to 1. Similarly to the expressions above, the probability *P*(*y* = 1) of the decision maker to select target *T*_1_ as decision *y* can be written as:

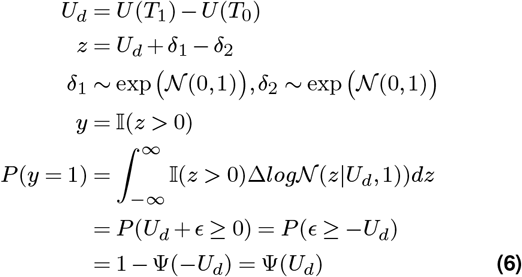

where 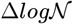 is the density function resulting from the difference of two log-normal distributions. A derivation for a numerical approximation of the cumulative density function of the difference between two (standard) log-normal distributions (Ψ) is given in the supplementary material.

The goal of these models is to estimate the structure of general preferences given only decisions over a subset of the whole possibility space. Our model is a special case in two regards: (1) Every decision is based on two alternatives only and (2) each subject is modeled individually. Since the RUMs do not make any assumptions about the features parameterizing and representing the utility function, we have to decide for its functional form. The two assumptions for this function were that (1) utilities only depend on amplitude, direction and change of direction of saccades and (2) these three features influence utilities independently of each other. The first assumption has the consequence, that preferences can only be recovered up to a common additive factor. The reason is, that this model only allows us to estimate the difference between the two utility values, which would be the same if both utility values had a constant part. This has the consequence, that we lose one degree of freedom within each part of the utility function. Thus one parameter can be set to an arbitrary value without loss of generality. The second assumption is known as *additive utility independence*. Accordingly, we define a potential saccadic target *T* in terms of a triplet of amplitude, direction and change of direction (amp*,ϕ,θ*). Now, to describe our utility function, we need to describe three independent utility functions, each describing the influence of a single feature. First, to choose a parameterization of the utilities depending on saccadic amplitude, we referred to the know phenomenon of saccadic proximity preference (Engel, 1971; Koch & Ullman, 1987). Previous research in motor control has demonstrated that endpoint variability of saccades grows linearly with saccadic amplitude, and that this property leads to a preference for shorter saccadic gaze shifts, if the precision of saccadic targeting is optimized (Harris & Wolpert, 1998). Therefore, utility in the present model was chosen to depend linearly on the amplitude of the gaze shift. The distance is normalized to the 0-1 interval by dividing saccadic amplitudes by the experimental maximum possible value, of 12 degree. Without loss of generality, this predominantly helps comparability and interpretability of the model’s results. This part of the utility function does not need a constant part because only utility differences are observed.

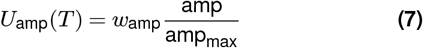

Second, many previous studies reported a preference for saccades in cardinal versus oblique directions, attributing this phenomenon to oculomotor biases (). But, simply rotating images revealed that the distribution of directions rotates with the image, providing evidence against the hypothesis that the initial distribution was due to oculomotor biases (Foulsham et al., 2008). Because our experiment had no stimuli or task information to promote cardinal eye movements, we wanted to keep the possible utility functions for the absolute direction in the visual field flexible. Therefore, we chose a piecewise linear function for *U_ϕ_*(*T*) with fixed turning points at 30°,90°,150°,210°,270°,330° and six variable function values at each of those turning points. Because angles are periodic around 360°, the interpolation below 30° or above 330° was done with the respective other counterpart. Since the model’s parameters can only be inferred from binary choices involving utility differences, *U_ϕ_* loses one degree of freedom. Without loss of generality, we fixed *w*_*ϕ*,1_ to 1.

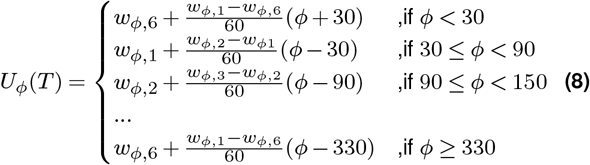

The third component of the utility function is the angle of a potential saccade relative to the previous gaze shift. The literature about the influence of past fixations contains numerous reports about return saccades being promoted (Luke et al., 2013; Luke et al., 2014; Wilming et al., 2013). For the current experiment a return saccade corresponds to a *θ* value of 0° or 360° degree. Furthermore, many studies treat the angular values as symmetric, therefore only calculating the smaller angle between two directions. Accordingly, our utility function is linear in the distance of *θ* to 180°. Similar to the amplitude, for reasons of parameter comparability, *θ* was normalized to the 0-1 interval:

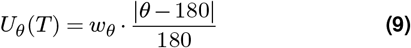

Having defined the above three component utility functions, the resulting overall utility of a saccade is modeled as their linear sum, see eq. 2.

### Inference of models’ parameters

The likelihood of a subject’s data is the product of individual trials’ likelihoods. This can be used to estimate parameters, either via gradient descent and maximum likelihood or using probabilistic programming in a Bayesian setting, which involves specifying prior distributions over parameters and allows computing posterior distributions over parameters. The overall data likelihood for a set of decisions *D*, consisting of tuples (*T*_1_,*T*_2_), where *T*_1_ always equals the chosen target, is:

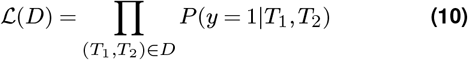

with the probability of target one to be chosen, from Equation 1. Parameters were estimated for each subject individually, using a graphical model, shown in Figure S5. Graphical models have the advantage that we do get an estimate of the posterior distribution of each parameter instead of only a point estimate of the most likely parameter configuration. Sampling was done in *Turing* with a *Hamiltonian Monte-Carlo* sampler(Duane et al., 1987; Ge et al., 2018). For each subject a chain of length 10000 with a burn-in of 1000 was sampled.

To compare the parameter posterior distributions of two subjects, we chose to compare the mean of the respective samples. Since all parameter distributions were unimodal and close to symmetric, the mean is a reasonable point to choose. To validate this assumption of the mean even further, parameters were also estimated using maximum likelihood estimation. This was done with the *Nelder-Mead* numerical optimization algorithm(Gao & Han, 2012; Virtanen et al., 2020). Indeed, deviations between the mean of the sampled chain and the maximum likelihood point estimate were minimal over all subjects and parameters (mean: 0.0137, SD: 0.0132, max: 0.0758, min: 0.000).

## Acknowledgements

This research was supported by the Hessian research priority program LOEWE within the project WhiteBox and the cluster projects “The Adaptive Mind” and “The Third Wave of AI” as part of the Excellence Program of the Hessian Ministry of Higher Education, Science, Research and Art.

## Supplementary Information

### S1 Derivation of the difference between two (standard) lognormal distributions

To infer the parameters of the utility function in a random utility model, we need the cumulative distribution function of the difference between the two noise distributions. Since we do not assume signal-dependant noise, both noise distributions are standard log-normal distributions. The difference between two log-normal distributions is known to have no close form solution, yet can be approximated via the following integral.

First some definitions, let *ϕ*(·) be the probability density of standard normal distribution 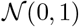, and Φ(·) its respective cumulative distribution. Moreover, we call *λ*(·) the probability density of the standard log-normal, and Λ(·) its respective cumulative function. Furthermore we call *Z* the random variable, which is the difference of the standard lognormal random variables, *X* and *Y*. Since *Z* is symmetric around 0, we only need to analyze for *z* > 0.

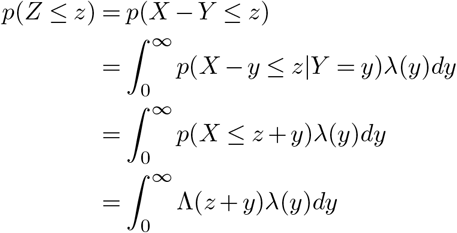

This last term can then be approximated with numerical integral solvers.

The rest of the derivation is analogous to random utility models with Gaussian uncertainties, leading to probit regression, from various textbooks(Murphy, 2021; Train, 2009).

### S2 Additional Figures and Tables

**Supplementary Figure S1.**
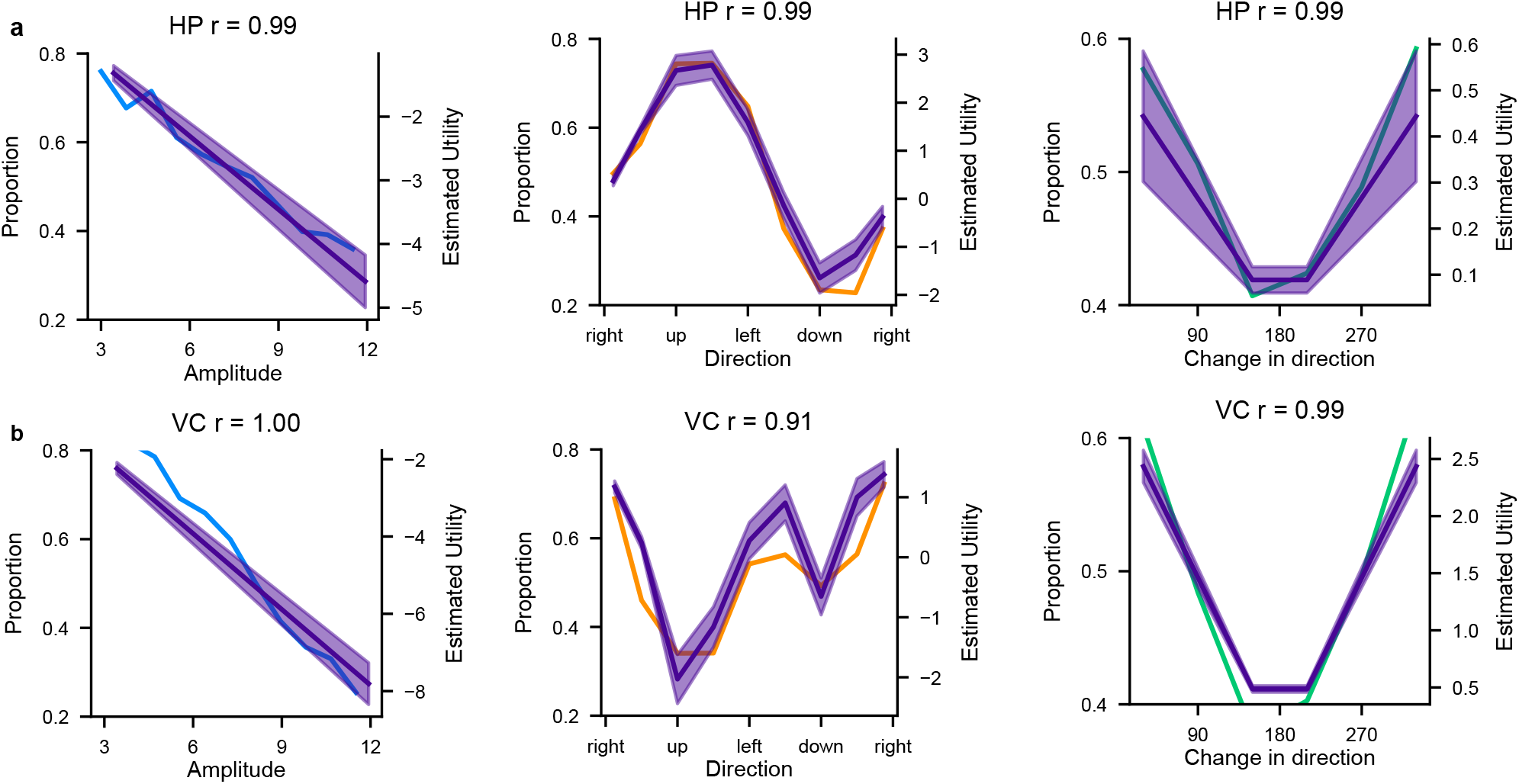
Comparison Model - Single subject preference. Comparisons for two subjects and all three features of their respective preferences and the predicted model utility values. The rows correspond to the subjects, while subplots in the same column correspond to the same feature. The colored line always stands for the proportion of targets chosen in a certain interval, while the purple line represents the modeled utility value for the center of that interval. The purple area indicates the mean of the respective parameter ± 2 standard deviations. The title contains the subject identifier, as well as the Pearson correlation between the two curves shown.

**Supplementary Figure S2.**
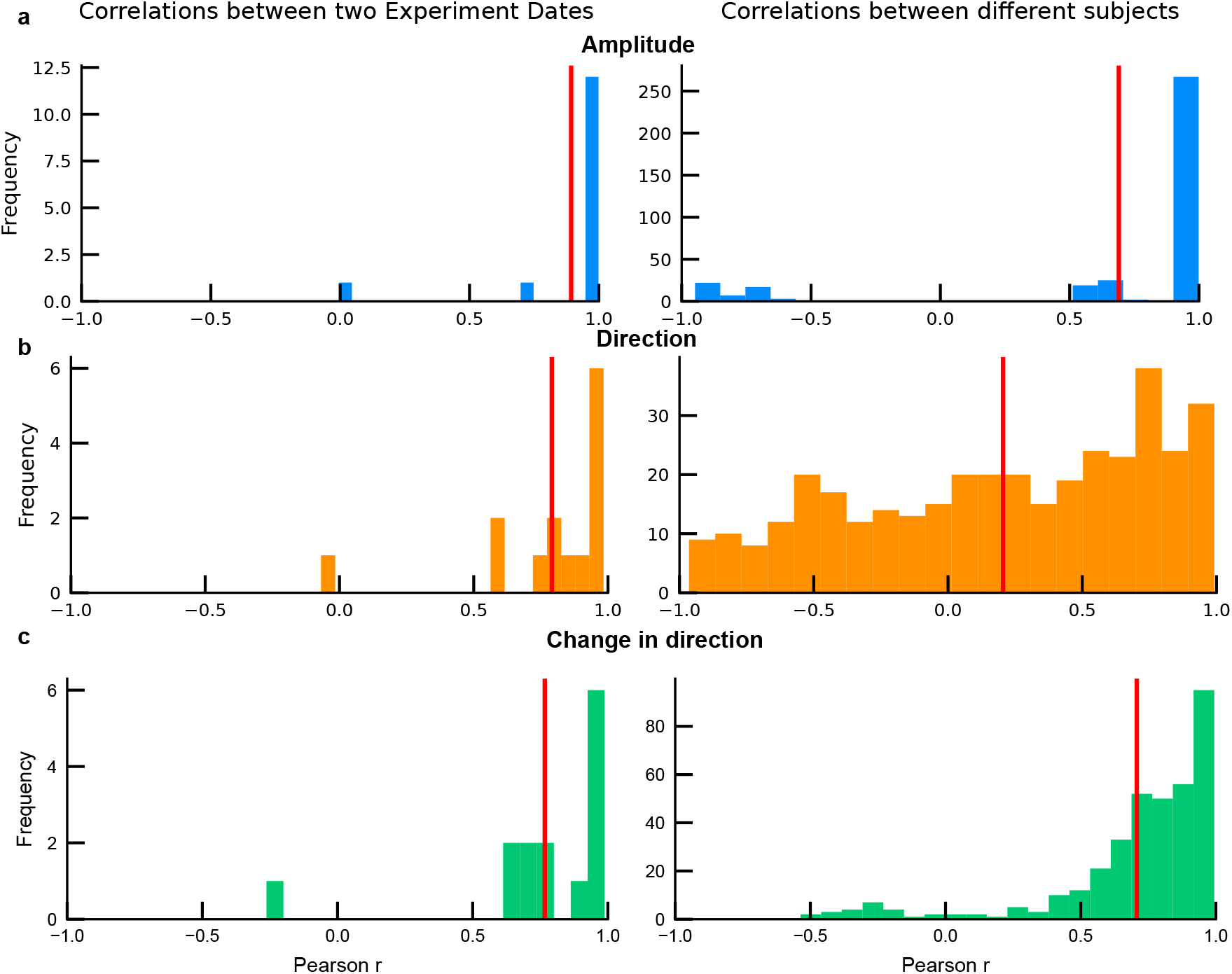
Correlations of proportions between experimental dates and between subjects. Left column: Distribution over correlation coefficients between the first and the second experiment date. Right column: Distribution over correlation coefficients between different subjects. Each row shows those distributions over one saccadic feature. The red line always indicates the mean over all subjects.

**Supplementary Figure S3.**
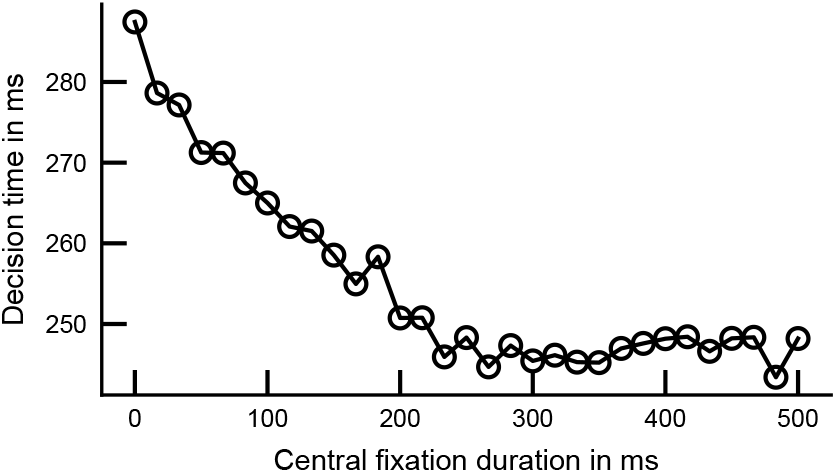
Decision time, dependant on the central fixation duration. The time in milliseconds subjects took to decide for one saccadic target and initiate the saccade, for all different fixation durations of the central target. The plot shows data averaged over all subjects.

**Supplementary Figure S4.**
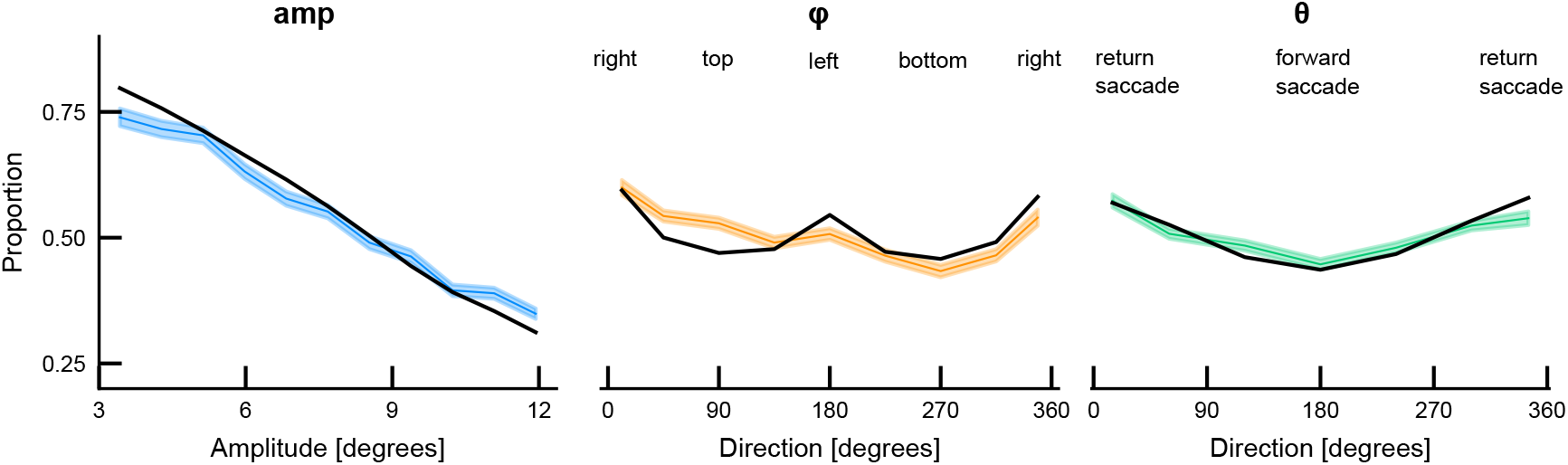
Comparison first 10 blocks - overall average. Comparison of the proportion of targets chosen between the first 10 blocks (colored) and the whole dataset (black). For the colored lines, the first 10 blocks (of the first experiment date) of every subject were included. That equals 10 minutes of experiment time per subject. Both lines show averages over all subjects. The shaded area around the curves is the standard error of the mean.

**Supplementary Figure S5.**
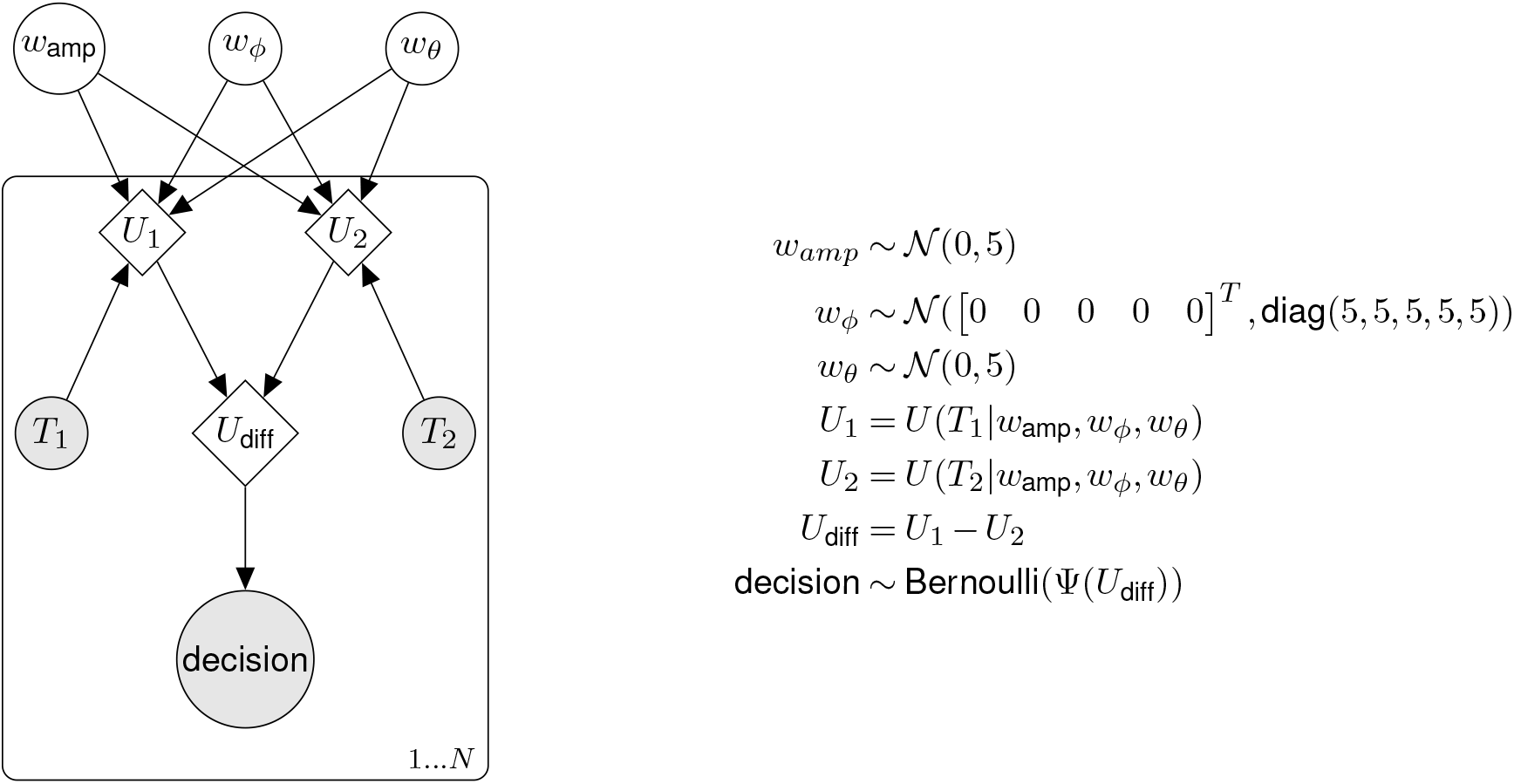
Bayesian network for estimating parameter posterior distributions. White nodes represent latent variables whereas shaded nodes represent observed quantities. Variables inside diamonds are deterministic, while the ones inside circles are random variables. The prior for all 7 parameters was a normal distribution with mean 0 and standard deviation of 5. Target *T*_1_ is always the target that the subject chose in a particular trial. The Bayesian network models every decision as a result of a Bernoulli random variable with *p* = *P*(*y* = 1) (from Equation 1) as the probability for the correct target.

**Supplementary Table S1.** Influence of experiment date and central fixation duration on preferences. All columns’ values are based on the distribution of mean squared errors between preference vectors over all participants. Both parts of the table calculate two preference vectors (all points visualized in Figure 2) for each participant and calculate the mean squared error between them. The upper part of the table describes the influence of the experiment date on the preferences. The lower part on the other hand, describes the difference between short (<250ms) and long (≥250ms) central fixation durations.

| Feature | mean | median | std | min | max |
| --- | --- | --- | --- | --- | --- |
| Difference between first and second experiment date |  |  |  |  |  |
| Amplitude | 0.0365 | 0.0281 | 0.0217 | 0.0163 | 0.1070 |
| Direction | 0.0671 | 0.0543 | 0.0519 | 0.0201 | 0.2279 |
| Change in direction | 0.0252 | 0.0223 | 0.0119 | 0.0085 | 0.0608 |
| Difference between short and long central fixation duration |  |  |  |  |  |
| Amplitude | 0.0290 | 0.0274 | 0.0084 | 0.0156 | 0.0480 |
| Direction | 0.0243 | 0.0246 | 0.0066 | 0.0099 | 0.0339 |
| Change in direction | 0.0189 | 0.0173 | 0.0070 | 0.0083 | 0.0340 |

## Bibliography

Anderson, A. J., Yadav, H., & Carpenter, R. H. S. (2008). Directional prediction by the saccadic system [Publisher: Elsevier]. Current biology, 18(8), 614–618.

Anderson, J. R. (1991). Is human cognition adaptive? Behavioral and Brain Sciences, 14(3), 471–485.

Bargary, G., Bosten, J. M., Goodbourn, P. T., Lawrance-Owen, A. J., Hogg, R. E., & Mollon, J. (2017). Individual differences in human eye movements: An oculomotor signature? Vision research, 141, 157–169.

Bays, P. M., & Husain, M. (2012). Active inhibition and memory promote exploration and search of natural scenes [Publisher: The Association for Research in Vision and Ophthalmology]. Journal of Vision, 12(8), 8–8. https://doi.org/10.1167/12.8.8

Bennett, P. J., & Pratt, J. (2001). The spatial distribution of inhibition of return. Psychological Science, 12(1), 76–80.

Borji, A., & Itti, L. (2012). State-of-the-art in visual attention modeling. IEEE transactions on pattern analysis and machine intelligence, 35(1), 185–207.

Borji, A., Sihite, D. N., & Itti, L. (2013). What stands out in a scene? a study of human explicit saliency judgment. Vision research, 91, 62–77.

De Haas, B., Iakovidis, A. L., Schwarzkopf, D. S., & Gegenfurtner, K. R. (2019). Individual differences in visual salience vary along semantic dimensions. Proceedings of the National Academy of Sciences, 116(24), 11687–11692.

Dodd, M. D., Van der Stigchel, S., & Hollingworth, A. (2009). Novelty is not always the best policy: Inhibition of return and facilitation of return as a function of visual task. Psychological Science, 20(3), 333–339.

Duane, S., Kennedy, A. D., Pendleton, B. J., & Roweth, D. (1987). Hybrid Monte Carlo. Physics Letters B, 195(2), 216–222. https://doi.org/10.1016/0370-2693(87)91197-X

Engel, F. (1971). Visual conspicuity, directed attention and retinal locus. Vision Research, 11(6), 563–575.

Findlay, J. M., & Gilchrist, I. D. (2003). Active vision: The psychology of looking and seeing. Oxford University Press.

Foulsham, T., Kingstone, A., & Underwood, G. (2008). Turning the world around: Patterns in saccade direction vary with picture orientation. Vision Research, 48(17), 1777–1790. https://doi.org/10.1016/j.visres.2008.05.018

Friedman, M., & Savage, L. J. (1952). The expected-utility hypothesis and the measurability of utility. Journal of Political Economy, 60(6), 463–474.

Gao, F., & Han, L. (2012). Implementing the Nelder-Mead simplex algorithm with adaptive parameters. Computational Optimization and Applications, 51(1), 259–277. https://doi.org/10.1007/s10589-010-9329-3

Ge, H., Xu, K., & Ghahramani, Z. (2018). Turing: Composable inference for probabilistic programming. In A. J. Storkey & F. Pérez-Cruz (Eds.), International Conference on Artificial Intelligence and Statistics, AISTATS 2018, 9-11 April 2018, Playa Blanca, Lanzarote, Canary Islands, Spain (pp. 1682–1690). PMLR. http://proceedings.mlr.press/v84/ge18b.html

Gershman, S. J., Horvitz, E. J., & Tenenbaum, J. B. (2015). Computational rationality: A converging paradigm for intelligence in brains, minds, and machines. Science, 349(6245), 273–278.

Gilchrist, I. D., & Harvey, M. (2000). Refixation frequency and memory mechanisms in visual search. Current Biology, 10(19), 1209–1212.

Gilchrist, I. D., & Harvey, M. (2006). Evidence for a systematic component within scan paths in visual search. Visual cognition, 14(4-8), 704–715.

Glimcher, P. W., & Fehr, E. (2013). Neuroeconomics: Decision making and the brain. Academic Press.

Gold, J. I., & Shadlen, M. N. (2007). The neural basis of decision making. Annu. Rev. Neurosci., 30, 535–574.

Gottlieb, J. (2012). Attention, learning, and the value of information. Neuron, 76(2), 281–295.

Harris, C. M., & Wolpert, D. M. (1998). Signal-dependent noise determines motor planning [Bandiera_abtest: a Cg_type: Nature Research Journals Number: 6695 Primary_-atype: Research Publisher: Nature Publishing Group]. Nature, 394(6695), 780–784. https://doi.org/10.1038/29528

Hayhoe, M., & Ballard, D. (2005). Eye movements in natural behavior. Trends in cognitive sciences, 9(4), 188–194.

Hayhoe, M., & Ballard, D. (2014). Modeling task control of eye movements. Current Biology, 24(13), R622–R628.

Henderson, J. M. (2003). Human gaze control during real-world scene perception. Trends in cognitive sciences, 7(11), 498–504.

Henderson, J. M., & Luke, S. G. (2014). Stable individual differences in saccadic eye movements during reading, pseudoreading, scene viewing, and scene search. Journal of Experimental Psychology: Human Perception and Performance, 40(4), 1390.

Hooge, I. T. C., Over, E. A., van Wezel, R. J., & Frens, M. A. (2005). Inhibition of return is not a foraging facilitator in saccadic search and free viewing. Vision research, 45(14), 1901–1908.

Hoppe, D., & Rothkopf, C. A. (2016). Learning rational temporal eye movement strategies. Proceedings of the National Academy of Sciences, 113(29), 8332–8337.

Hoppe, D., & Rothkopf, C. A. (2019). Multi-step planning of eye movements in visual search. Scientific reports, 9(1), 1–12.

Horowitz, T. S., & Wolfe, J. M. (1998). Visual search has no memory. Nature, 394(6693), 575–577.

Klein, R. M., & MacInnes, W. J. (1999). Inhibition of return is a foraging facilitator in visual search. Psychological science, 10(4), 346–352.

Koch, C., & Ullman, S. (1987). Shifts in selective visual attention: Towards the underlying neural circuitry. In Matters of intelligence (pp. 115–141). Springer.

Körding, K. P., Fukunaga, I., Howard, I. S., Ingram, J. N., & Wolpert, D. M. (2004). A Neuroeconomics Approach to Inferring Utility Functions in Sensorimotor Control (James Ashe, Ed.). PLoS Biology, 2(10), e330. https://doi.org/10.1371/journal.pbio.0020330

Land, M., & Tatler, B. (2009). Looking and acting: Vision and eye movements in natural behaviour. Oxford University Press.

Lisi, M., Solomon, J. A., & Morgan, M. J. (2019). Gain control of saccadic eye movements is probabilistic. Proceedings of the National Academy of Sciences, 116(32), 16137–16142.

Luke, S. G., Schmidt, J., & Henderson, J. M. (2013). Temporal oculomotor inhibition of return and spatial facilitation of return in a visual encoding task [Publisher: Frontiers]. Frontiers in Psychology, 0. https://doi.org/10.3389/fpsyg.2013.00400

Luke, S. G., Smith, T. J., Schmidt, J., & Henderson, J. M. (2014). Dissociating temporal inhibition of return and saccadic momentum across multiple eye-movement tasks [Publisher: The Association for Research in Vision and Ophthalmology]. Journal of Vision, 14(14), 9–9.

McFadden, D., et al. (1973). Conditional logit analysis of qualitative choice behavior.

Murphy, K. P. (2021). Machine learning: A probabilistic perspective [OCLC: 1255636989]. MIT Press.

Navalpakkam, V., Koch, C., Rangel, A., & Perona, P. (2010). Optimal reward harvesting in complex perceptual environments. Proceedings of the National Academy of Sciences, 107(11), 5232–5237.

Peterson, M. S., Kramer, A. F., Wang, R. F., Irwin, D. E., & McCarley, J. S. (2001). Visual search has memory. Psychological Science, 12(4), 287–292.

Petitet, P., Attaallah, B., Manohar, S. G., & Husain, M. (2021). The computational cost of active information sampling before decision-making under uncertainty. Nature Human Behaviour, 5(7), 935–946.

Risko, E. F., Anderson, N. C., Lanthier, S., & Kingstone, A. (2012). Curious eyes: Individual differences in personality predict eye movement behavior in scene-viewing. Cognition, 122(1), 86–90.

Rothkopf, C. A., Ballard, D. H., & Hayhoe, M. M. (2007). Task and context determine where you look. Journal of vision, 7(14), 16–16.

Schütz, A. C., Trommershäuser, J., & Gegenfurtner, K. R. (2012). Dynamic integration of information about salience and value for saccadic eye movements. Proceedings of the National Academy of Sciences, 109(19), 7547–7552.

Schwarz, G. (1978). Estimating the Dimension of a Model. The Annals of Statistics, 6(2). https://doi.org/10.1214/aos/1176344136

Simon, H. A. (1955). A behavioral model of rational choice. The quarterly journal of economics, 69(1), 99–118.

Smith, T. J., & Henderson, J. M. (2009). Facilitation of return during scene viewing [Publisher: Taylor & Francis]. Visual Cognition, 17(6-7), 1083–1108.

Tatler, B. W., Baddeley, R. J., & Vincent, B. T. (2006). The long and the short of it: Spatial statistics at fixation vary with saccade amplitude and task. Vision Research, 46(12), 1857–1862. https://doi.org/10.1016/j.visres.2005.12.005

Tatler, B. W., & Vincent, B. T. (2009). The prominence of behavioural biases in eye guidance. Visual Cognition, 17(6-7), 1029–1054. https://doi.org/10.1080/13506280902764539

Thorndike, R. L. (1953). Who belongs in the family. Psychometrika.

Thurstone, L. L. (1927a). A law of comparative judgment. [Publisher: Psychological Review Company]. Psychological review, 34(4), 273.

Thurstone, L. L. (1927b). Psychophysical analysis [Publisher: JSTOR]. The American journal of psychology, 38(3), 368–389.

Todorov, E., & Jordan, M. I. (2002). Optimal feedback control as a theory of motor coordination. Nature neuroscience, 5(11), 1226–1235.

Torralba, A., Oliva, A., Castelhano, M. S., & Henderson, J. M. (2006). Contextual guidance of eye movements and attention in real-world scenes: The role of global features in object search. Psychological review, 113(4), 766.

Train, K. (2009). Discrete choice methods with simulation [OCLC: 1104459815]. Cambridge University Press. Retrieved September 7, 2021, from http://proxy.uqtr.ca/login.cgi?action=login&u=uqtr&db=ebsco&ezurl=http://search.ebscohost.com/login.aspx?direct=true&scope=site&db=nlebk&AN=304795

Virtanen, P., Gommers, R., Oliphant, T. E., Haberland, M., Reddy, T., Cournapeau, D., Burovski, E., Peterson, P., Weckesser, W., Bright, J., van der Walt, S. J., Brett, M., Wilson, J., Millman, K. J., Mayorov, N., Nelson, A. R. J., Jones, E., Kern, R., Larson, E., … Vázquez-Baeza, Y. (2020). SciPy 1.0: Fundamental algorithms for scientific computing in Python. Nature Methods, 17(3), 261–272. https://doi.org/10.1038/s41592-019-0686-2

Von Neumann, J., & Morgenstern, O. (1944). Theory of games and economic behavior. Princeton University Press.

Wilming, N., Harst, S., Schmidt, N., & König, P. (2013). Saccadic Momentum and Facilitation of Return Saccades Contribute to an Optimal Foraging Strategy (O. Sporns, Ed.). PLoS Computational Biology, 9(1), e1002871. https://doi.org/10.1371/journal.pcbi.1002871

Yarbus, A. L. (1967). Eye movements and vision. New York: Plenum Press.

